# In Vitro Assessment of Chemically Modified Probiotic Bacteria as Receptor-Mimic Decoys for Cholera Toxin B Subunit

**DOI:** 10.64898/2026.09.30.755077

**Authors:** Ilaria Mainero Rocca, Nicolai Stuhr-Hansen, Ola Blixt, Steffen Goletz

## Abstract

Gastrointestinal infections remain a major cause of morbidity and mortality worldwide. Conventional antibiotic treatment can be ineffective or inappropriate in some settings, particularly in the context of increasing antimicrobial resistance. In many enteric infections, an early step in pathogenesis involves binding of pathogens or their toxins to oligosaccharide receptors on host-cell surfaces, making these interactions attractive targets for antimicrobial strategies that do not rely on conventional antibiotics. Here, we developed a non-recombinant, microbiome-based chemical engineering approach in which cholesteryl-peptides serve as flexible chemical scaffolds for displaying GM1 receptor mimics on probiotic bacteria. The GM1-cholesteryl-peptides generated surface-accessible GM1 on both the Gram-positive probiotic Lactobacillus reuteri DSM 17938 and the Gram-negative probiotic Escherichia coli Nissle 1917. The resulting GM1-decorated bacteria showed concentration-dependent binding and sequestration of cholera toxin B subunit (CTB) in vitro and reduced CTB binding to Vero cells. These findings establish an in vitro proof-of-concept for a modular, non-recombinant bacterial decoy platform and may provide a basis for the future development of receptor-mimic strategies against toxin-mediated gastrointestinal infections.

## 1. Introduction

Infectious diseases, including gastrointestinal (GI) infections, remain a major public-health burden worldwide [1,2]. Cholera continues to contribute substantially to global morbidity and mortality [2]. A major challenge in the treatment of infectious diseases is the emergence of multidrug-resistant strains [3,4]. The development of new antibiotics requires considerable time and investment and may not keep pace with the rapid emergence of resistance [5]. There is therefore an urgent need for alternative strategies that do not rely exclusively on conventional antibiotics. Many interactions involved in GI infection are mediated by glycans and glycan-binding receptors expressed by microbial pathogens and host cells [6]. Accordingly, antimicrobial approaches that block glycan-receptor interactions between GI cells and pathogens or toxins have been explored. For example, Synsorb-Pk is a nanostructured toxin-binding system in which the synthetic receptor Gb3, the carbohydrate determinant for Shiga toxin (Stx) binding [7], was covalently coupled to inert silica particles [8]. Although Synsorb-Pk bound Stx and inhibited its cytotoxicity in vitro [9], oral administration did not reduce the severity of diarrhea-associated hemolytic uremic syndrome [10]. Similarly, Das and colleagues developed poly(lactic-co-glycolic acid)-based nanoparticles coated with polyethylene glycol and GM1 oligosaccharide, the principal host receptor for cholera toxin (CT), as decoys for the enterotoxin [11]. Although nanoparticles have potential as anti-infective agents, their metabolism, clearance, and toxicity require further evaluation before clinical translation [12]. A more recent strategy involves recombinant probiotics engineered to prevent or limit infections, including cholera [13–18]. Clinical studies have also supported the use of probiotics such as Lactobacillus reuteri to reduce the incidence or duration of diarrheal disease [19–24]. The Gram-negative Escherichia coli Nissle 1917 (EcN) is a well-established probiotic strain [25] that has also been used as a chassis for engineered probiotic approaches, including strategies to compete with other bacteria [14], target pathogens associated with GI infections such as Vibrio cholerae [15], or neutralize CT [16–18]. Such probiotic-engineering approaches have been reviewed extensively [26] and, when recombinant, are subject to regulatory considerations associated with genetically modified microorganisms [27,29]. A non-recombinant platform may avoid some GMO-specific considerations, although non-recombinant status does not in itself establish safety. In this context, previous studies have identified probiotic bacteria capable of binding or removing cholera toxin, supporting the potential development of non-antibiotic, microbiome-based toxin-binding strategies [28,30]. We therefore set out to develop a non-recombinant, chemically modified probiotic platform based on neo-glycopeptide modules bearing a GM1 oligosaccharide as a molecular mimic for CTB recognition. Here, we show that GM1 ganglioside and GM1-neoglycopeptides generate CTB-accessible GM1 signals on both a Gram-positive and a Gram-negative probiotic strain, and that the modified bacterial scaffolds reduce binding of soluble CTB to GM1-expressing Vero cells in vitro.

## 2. Results and Discussion

### 2.1 Synthesis of cholesterylated neo-glycopeptide modules

To decorate bacterial surfaces with functional molecules capable of binding the GM1-recognizing cholera toxin B subunit (CTB), we adapted our previously reported glycocalyxification strategy [31] to synthesize cholesteryl-peptides bearing functional groups for the facile introduction of azido-glycans, followed by click conjugation with an azide-functionalized GM1 oligosaccharide [32] (Scheme 1). Related modules had previously been shown to insert into artificial lipid membranes, including giant unilamellar vesicles (GUVs), as well as into natural membranes such as those of red blood cells, while remaining accessible to lectins and cellular receptors [33,34]. We therefore hypothesized that analogous compounds could associate with bacterial envelopes. The GM1-cholesterylated neo-glycopeptide modules were synthesized by solid-phase peptide synthesis using our previously reported Fmoc-cholesterol building block (Fmoc-Ala-cholesteryltriazyl-OH), Fmoc-OEG2-OH (Fmoc-O2Oc-OH), and Fmoc-Pra-OH (Fmoc-propargyllysine) as standard Fmoc amino-acid derivatives, with a propargyl anchor separated from a biotin recognition moiety by a PEG spacer [31] (Scheme 1). The molecular design incorporated either a short Lys-Lys dipeptide, providing a positively charged module, or a Glu-Lys dipeptide, providing an overall neutral module, to mitigate the hydrophobicity of the cholesteryl moiety and improve aqueous handling. Biotin was introduced at the N terminus to enable subsequent detection. After TFA-based cleavage and HPLC purification (>98% purity), HRMS confirmed the identities of the biotinylated Lys-Lys and Glu-Lys propargylated cholesteryl-peptides 1 and 2. Compounds 1 and 2 were subsequently reacted with GM1 azide under CuI-catalyzed click-chemistry conditions to yield GM1-Lys-Lys-cholesteryl-peptide (GM1LL; compound 3) and GM1-Glu-Lys-cholesteryl-peptide (GM1GL; compound 4), respectively (Scheme 1); these two GM1-functionalized cholesteryl-peptides are collectively referred to below as GM1chol. The GM1 azide used for conjugation was obtained through the Consortium for Functional Glycomics (CFG). Detailed synthetic procedures and reagent provenance are provided in Supporting Information, Appendix I.

**Scheme 1.**
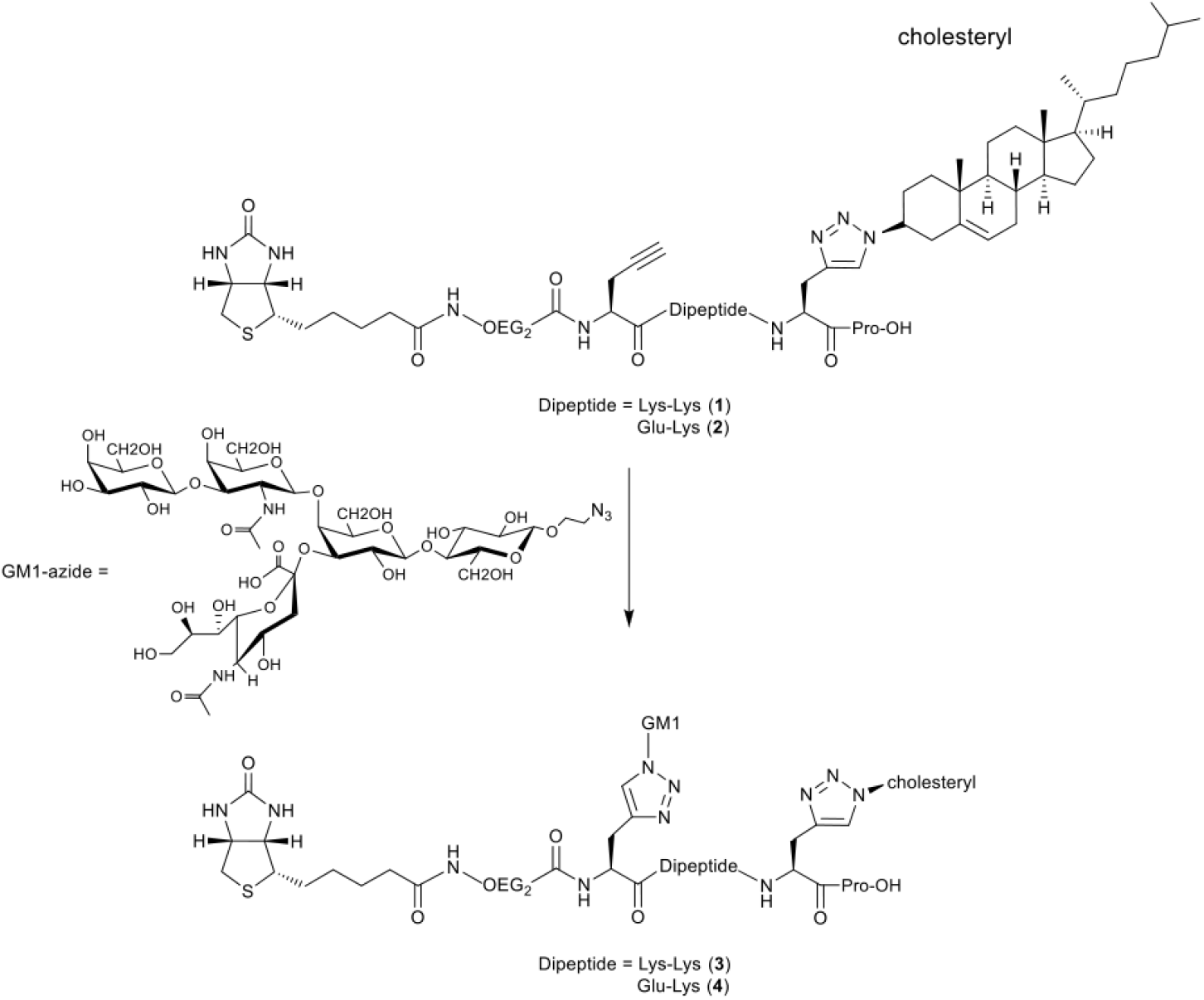
Synthesis of biotin-propargyl-functionalized cholesteryl-peptides. Biotin-propargyl-functionalized cholesteryl-peptides 1 and 2 were obtained by SPPS on 2-chlorotrityl polystyrene resin preloaded with proline (H-Pro-2Clrt). Following biotinylation, GM1-functionalized cholesteryl-peptides 3 and 4 were generated by click chemistry using a CuI-based catalyst. Reaction conditions: (i) SPPS on 2-chlorotrityl resin: proline (1 equiv.), Fmoc-Ala(Cholesteryl)-OH (4 equiv.), Fmoc-propargyl-lysine (4 equiv.), Fmoc-O2Oc-OH (4 equiv.), HBTU (4 equiv.), DIPEA, NMP, 30 min, room temperature (rt). (ii) Biotinylation: 1 and 2 (0.1 equiv.), biotin (0.5 equiv.), HBTU (0.5 equiv.), DIPEA:NMP (1:6), 30 min, rt. (iii) Click chemistry: 1 (0.3 equiv.) or 2 (0.3 equiv.), GM1 azide (0.3 equiv.), CuI (1 mol%), DIPEA:MeOH (1:9), 16 h, rt.

### 2.2 Assembly, detection and CTB sequestration by GM1-neoglycopeptide-loaded probiotics

Lactobacillus reuteri DSM 17938 (LR) and Escherichia coli Nissle 1917 (EcN) were selected as probiotic bacterial scaffolds for generating CTB-binding surfaces. To determine whether incubation with GM1-cholesteryl-peptides (GM1chol) generated surface-accessible GM1, the strains were incubated with the modules and subsequently assessed for CTB binding. Loading conditions were first optimized using natural GM1 ganglioside as a benchmark to determine whether similar conditions could be applied to the cholesteryl-peptides (Appendix II, Supplementary Figures 1 and 2). CTB binding increased with GM1 loading in both LR and EcN. No clear relationship between bacterial density and toxin-binding capacity was observed; however, lower OD600 values appeared to require less GM1 loading to approach maximal CTB binding (Appendix II, Supplementary Figures 1 and 2). A complementary experiment likewise detected CTB-accessible GM1 and GM1chol after incubation of LR and EcN with the corresponding molecules (Appendix II, Supplementary Figure 3). Based on these optimization experiments, and to maintain sufficient bacterial material throughout the assay, OD600 = 2 was selected for subsequent experiments. After overnight culture, LR and EcN were resuspended in loading buffer at OD600 = 2 and incubated with increasing concentrations of GM1 ganglioside (GM1gs), GM1chol, or propargyl-cholesteryl-peptides lacking the GM1 sugar moiety as controls. After removal of unbound loading molecules, the bacteria were incubated with CTB-HRP and analyzed by ELISA (Figure 1A). Both LR and EcN showed concentration-dependent CTB-HRP binding after loading with GM1gs, GM1GL, or GM1LL compared with control-loaded bacteria (Figure 1B(i–iii), C(i–iii)). Previous studies have demonstrated association of GM1 ganglioside or cholesterol-containing molecules with phospholipid bilayers [31,35]; our ELISA data are consistent with the presence of CTB-accessible GM1 on the bacterial surface after loading. The signal observed with LR is particularly notable given its thick peptidoglycan layer, although the molecular basis and precise localization of the loaded molecules remain to be determined. Approximately ten-fold higher concentrations of GM1GL and GM1LL than of GM1gs were required to approach maximal CTB binding. This difference may reflect uncertainty in the concentrations of the synthetic products and/or differences in the accessibility or orientation of the GM1 moiety at the bacterial surface. Importantly, CTB binding is an indirect readout that depends on both the amount of surface-associated GM1 and its accessibility and orientation; it therefore does not provide a direct quantitative measure of membrane incorporation. Taken together, these results are consistent with successful generation of surface-accessible GM1 on LR and EcN and its specific recognition by CTB.

**Figure 1.**
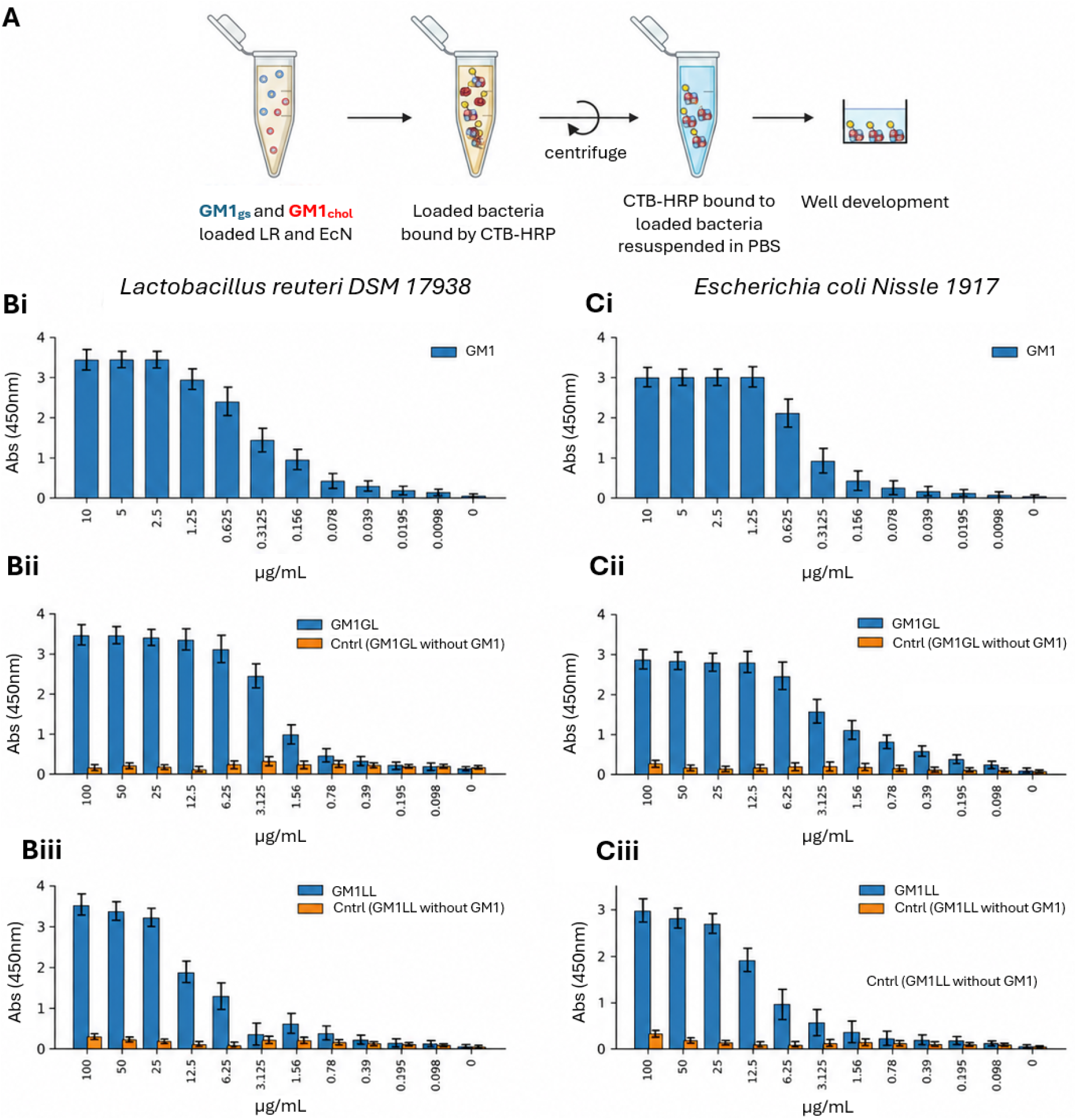
Concentration-dependent CTB-HRP recognition of surface-accessible GM1 on LR and EcN after loading with GM1 lipids. A) Schematic representation of the assay. LR and EcN at OD600 = 2 were loaded with increasing concentrations of GM1gs (0–10 µg/mL), GM1GL, GM1LL, or the corresponding cholesteryl controls (0–100 µg/mL), followed by CTB-HRP detection by ELISA. B(i–iii) CTB-HRP binding to LR loaded with GM1gs (i), GM1GL (ii), or GM1LL (iii). C(i–iii) Corresponding CTB-HRP measurements for EcN. Data are shown as mean ± SD of the experimental repeats (n = 3 for GM1gs and GM1GL; n = 2 for GM1LL).

After establishing CTB recognition by the loaded bacteria, we quantified depletion of CTB-HRP from the supernatant using an ELISA-based assay. A fixed concentration of CTB-HRP was incubated with bacteria loaded with increasing concentrations of GM1gs, GM1GL, GM1LL, or the corresponding cholesteryl controls. Unbound CTB-HRP was transferred to GM1-coated plates and quantified against a CTB-HRP standard curve (Appendix II, Supplementary Figure 4A). GM1GL and GM1LL produced a concentration-dependent reduction in free CTB-HRP, whereas the cholesteryl controls lacking GM1 showed no comparable effect (Appendix II, Supplementary Figure 4B-E). The lowest loading concentration associated with >95% depletion of CTB-HRP was approximately 10 µg/mL GM1GL for both LR and EcN, 25 µg/mL GM1LL for EcN, and 50 µg/mL GM1LL for LR. GM1gs produced a comparable effect at lower loading concentrations (2.5 µg/mL for EcN and 5 µg/mL for LR).

We next tested whether CTB sequestration by the loaded bacteria reduced CTB binding to Vero cells. A fixed concentration of Alexa Fluor 555-labelled CTB (CTB-555) was pre-incubated with LR or EcN loaded with increasing concentrations of GM1gs, GM1GL, GM1LL, or the corresponding cholesteryl controls. After centrifugation, the supernatants containing residual unbound CTB-555 were transferred to Vero cells and cell-associated fluorescence was quantified by flow cytometry (Figure 2A). Figure 2B and C show the combined quantitative data from two separately recorded experimental datasets. Within each dataset, mean PE-A fluorescence was normalized to the corresponding 0 µg/mL bacterial-loading condition (100% CTB-555 binding) before the datasets were combined. A concentration-dependent reduction in Vero-cell-associated CTB-555 fluorescence was observed after pre-incubation with GM1gs-, GM1GL-, and GM1LL-loaded LR and EcN, whereas the cholesteryl controls remained close to the unloaded condition. Nonlinear regression of the combined normalized data yielded apparent IC50 values of 0.54 µg/mL (95% CI, 0.40–0.69) for GM1gs, 4.5 µg/mL (3.2–5.7) for GM1LL, and 1.8 µg/mL (1.5–2.1) for GM1GL in LR; the corresponding values for EcN were 0.59 µg/mL (0.28–0.91), 3.0 µg/mL (2.4–3.7), and 1.4 µg/mL (0.67–2.1), respectively. The flow-cytometry gating workflow is described in Methods Section 3.5 and illustrated in Appendix II, Supplementary Figure 5.

**Figure 2.**
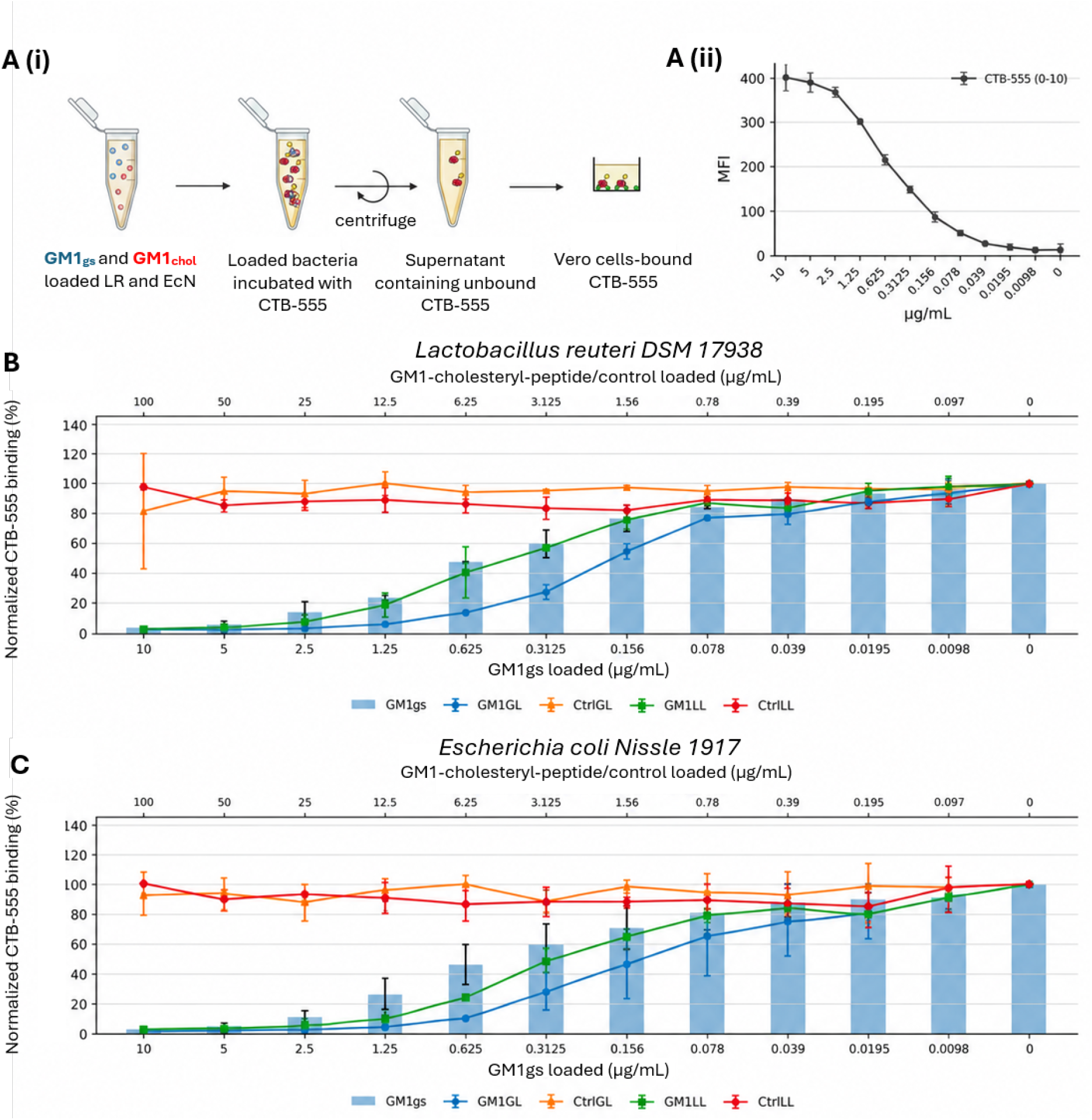
Reduction of CTB binding to Vero cells following pre-incubation with GM1-lipid-decorated LR and EcN, evaluated by flow cytometry. A(i) Schematic representation of the assay. LR and EcN at OD600 = 2, loaded with GM1gs (0–10 µg/mL) or GM1-cholesteryl-peptides (GM1chol; 0–100 µg/mL), were incubated with Alexa Fluor 555-labelled CTB (CTB-555). After centrifugation, supernatants containing unbound CTB-555 were transferred to Vero cells. A(ii) CTB-555 binding to Vero cells across a 0–10 µg/mL titration. B,C) Concentration-dependent reduction of CTB-555 binding to Vero cells after pre-incubation with LR (B) or EcN (C) loaded with GM1gs, GM1GL, GM1LL, or the corresponding cholesteryl controls. GM1gs concentrations are shown on the lower x-axis and GM1-cholesteryl-peptide/control concentrations on the upper x-axis. Within each of two separately recorded experimental datasets, mean PE-A fluorescence was normalized to the corresponding 0 µg/mL loading condition (100% CTB-555 binding). Data are shown as mean ± SD (n = 2 separately recorded experimental datasets). A representative gating strategy is provided in Appendix II, Supplementary Figure 5.

These results support concentration-dependent sequestration of CTB and reduced CTB binding to Vero cells by GM1-decorated bacterial scaffolds. However, CTB contains the GM1-binding B subunit but lacks the enzymatically active A subunit; therefore, the present experiments do not demonstrate neutralization of intact AB5 cholera toxin or protection from toxin-induced cellular effects. The findings should thus be interpreted as an in vitro proof-of-concept for CTB sequestration and reduced receptor binding, rather than as evidence of therapeutic efficacy against cholera.

Compared with the recombinant receptor-mimic E. coli strategy reported by Focareta et al. [18], the present platform introduces the toxin-binding glycan by non-recombinant chemical surface modification rather than by genetically encoded biosynthesis. This design provides a modular platform in which the displayed glycan or membrane anchor could, in principle, be varied without genetically re-engineering the bacterial strain, potentially facilitating preparation of alternative receptor-mimic surfaces. The non-recombinant format may also avoid some regulatory considerations that are specific to genetically modified microorganisms. Conversely, unlike genetically encoded systems, the chemical surface modification is not self-renewing: surface-associated molecules may be diluted or lost during bacterial growth and division, and their retention, orientation, and accessibility may vary with bacterial membrane architecture and environmental conditions. These trade-offs will require evaluation in future in vivo studies.

### 2.3 Stability of CTB-binding capacity following stress treatments

To assess whether CTB-binding capacity associated with the chemically modified bacterial surface was retained after stresses relevant to oral delivery and storage, GM1-loaded LR and EcN were subjected to acidic pH, heat, freezing/thawing, and lyophilization before CTB-HRP detection (Figure 3A). GM1gs was used at 1 µg/mL, whereas GM1GL, GM1LL, and the related controls were used at 10 µg/mL. After exposure to HCl at pH 2 for 1 h at 37°C, the mean CTB-binding signal decreased by approximately 16% and 22% for GM1gs-loaded LR and EcN, respectively, and by approximately 39% and 10% for GM1LL-loaded LR and EcN, whereas GM1GL-dependent CTB recognition was largely retained (Figure 3B). Because CTB binding reflects both the amount and the accessibility/orientation of surface-associated GM1, these measurements should be interpreted as retention of CTB recognition rather than as direct quantification of membrane incorporation. Heat treatment at 70°C for 15 min produced the largest reduction for GM1LL-loaded LR (approximately 66%), with smaller reductions for GM1LL-loaded EcN (approximately 34%) and GM1gs-loaded EcN (approximately 31%); GM1gs-loaded LR and GM1GL-loaded bacteria showed smaller changes (Figure 3C). Following freezing/thawing, CTB-binding signals were largely retained for GM1gs and GM1GL, whereas GM1LL-loaded LR showed an approximately 23% decrease (Figure 3D). After lyophilization, the largest decreases were observed for GM1LL-loaded LR (approximately 33%) and GM1gs-loaded EcN (approximately 24%), while the remaining conditions showed comparatively small changes (Figure 3E). These observations are descriptive because the available dataset comprised three experimental repeats for GM1gs and GM1GL and two for GM1LL; technical duplicate measurements were averaged within each repeat, and no inferential statistical testing was applied. Importantly, these experiments assessed retention of CTB-binding capacity and did not measure bacterial viability. In particular, exposure to 70°C for 15 min would be expected to compromise viability even if surface-associated GM1 remained detectable. Thus, the present data support stability of the toxin-binding surface signal, not preservation of probiotic viability or function. The persistence of CTB binding after treatments that may reduce viability also raises the possibility that viable bacteria may not be strictly required for a future toxin-binding scaffold, although this possibility was not tested here. Finally, exposure to pH 2 for 1 h represents only an initial in vitro approximation of gastric stress. The gastrointestinal environment also contains bile salts, digestive enzymes, mucus, competing lipids and proteins, and dynamic transit conditions, all of which may affect retention and accessibility of the displayed GM1. These factors, together with bacterial viability, intestinal retention, and toxin-dependent pathology, will need to be evaluated in physiologically relevant models before therapeutic application can be assessed. Lyophilization or pH-responsive encapsulation may be considered in future formulation studies [36]. Overall, the present experiments demonstrate partial retention of CTB-binding capacity after the tested stresses and support further evaluation of this non-recombinant bacterial decoy platform.

**Figure 3.**
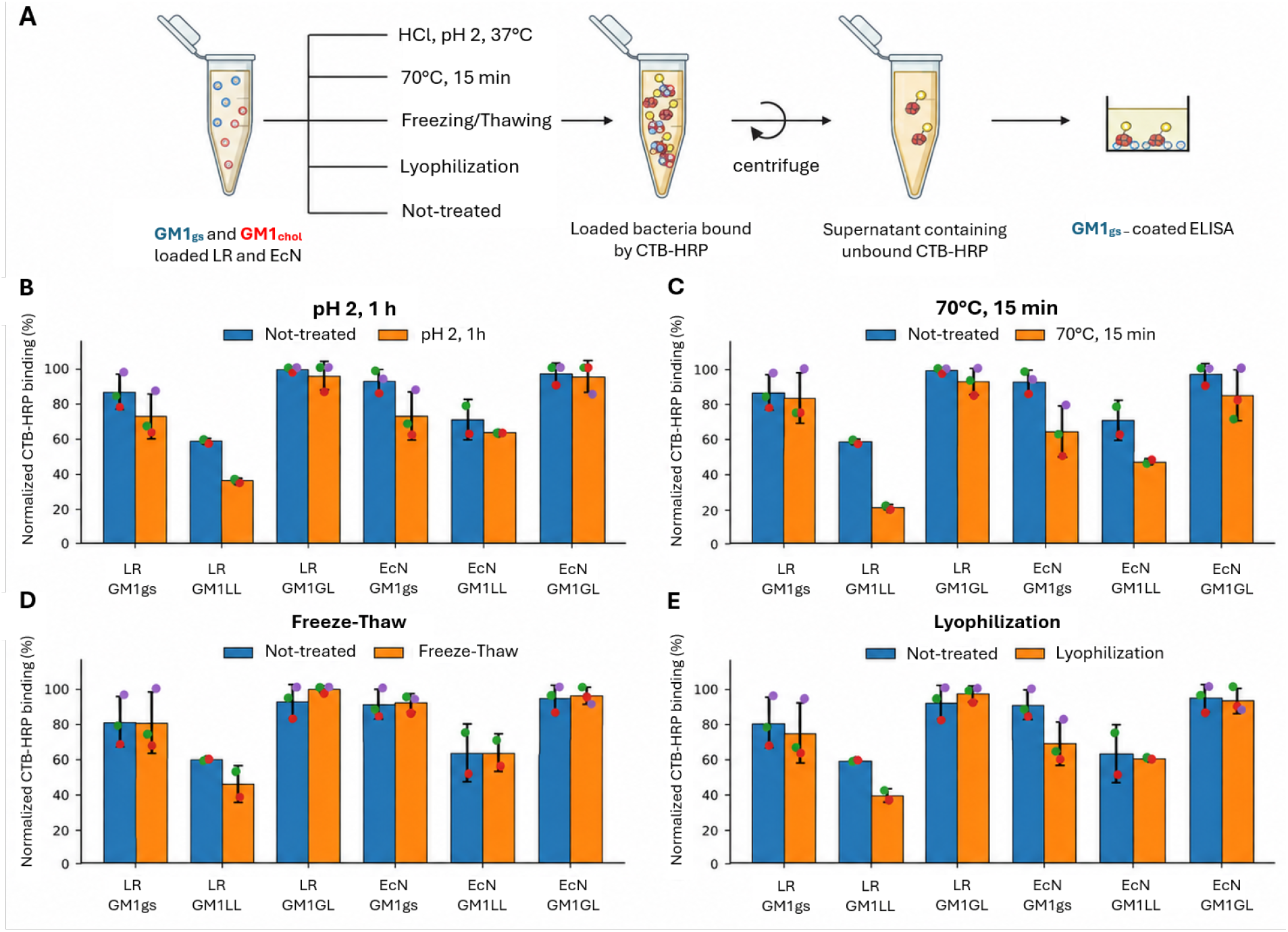
In vitro stability of CTB-binding capacity following stress treatments. A) Schematic representation of the assay. LR and EcN at OD600 = 2, loaded with either GM1gs (1 µg/mL) or GM1chol (10 µg/mL), were subjected to pH 2, 70°C for 15 min, freezing/thawing, lyophilization, or no treatment before probing with HRP-labelled CTB. After centrifugation and washing, bacterial suspensions were assayed for CTB-HRP binding in an ELISA plate. B-E) Retained CTB-binding signal after pH 2 exposure (B), heat treatment (C), freezing/thawing (D), and lyophilization (E). These measurements assess CTB recognition of surface-accessible GM1 and do not measure bacterial viability or directly quantify membrane incorporation. Technical duplicate measurements were averaged within each experimental repeat. Data are shown as mean ± SD of the experimental repeats, with individual repeat means overlaid (n = 3 for GM1gs and GM1GL; n = 2 for GM1LL). Owing to the limited number of experimental repeats, no inferential statistical testing was performed.

## 3. Methods

### 3.1 Preparation of loaded probiotics

Lactobacillus reuteri DSM 17938 and E. coli Nissle 1917 were kindly provided by Prof. Susanne Brix (Technical University of Denmark) and Assistant Prof. Nathan Crook (North Carolina State University), respectively. L. reuteri DSM 17938 was grown overnight at 37°C in MRS broth (Sigma) containing 1% Tween 80 in 50 mL Falcon tubes under anaerobic conditions, using either glycerol-stock inoculum or colonies from MRS agar plates. E. coli Nissle 1917 was grown overnight at 37°C and 220 rpm in LB broth (Sigma) in 50 mL Falcon tubes, using either glycerol-stock inoculum or colonies from LB agar plates. The following day, cultures were centrifuged at 4000 × g for 10 min, resuspended in PBS, centrifuged again, and transferred to BSM (Sigma), a suboptimal medium that limits bacterial growth, for overnight incubation at 37°C. Bacteria were then washed three times with PBS (each wash consisting of centrifugation at 4000 × g for 10 min followed by resuspension in PBS) and adjusted to the desired OD600. Suspensions were distributed into deep V-bottom polypropylene plates (Fisher Scientific) at 100 µL/well and centrifuged at 1000 × g for 10 min. After removal of the PBS, bacterial pellets were loaded with 100 µL/well of GM1 (Sigma; 0–10 µg/mL) or GM1-cholesteryl-peptides and the corresponding structural controls (0– 100 µg/mL) in PBS:BSM (1:1) loading buffer and incubated overnight at 37°C and 220 rpm. The cells were subsequently washed three times with PBS containing 0.05% Tween 20 to remove unbound loading molecules.

### 3.2 GM1-CTB enzyme-linked immunosorbent assay (ELISA)

Monosialoganglioside GM1 (Sigma) was diluted in PBS (pH 7.2), and 50 µL was added to each well of a Maxisorp ELISA plate and adsorbed overnight at 4°C. Coated wells were washed three times with 200 µL PBS containing 0.05% Tween 20; the same washing procedure was used after each subsequent incubation step. Wells were blocked for 30 min with 200 µL PBS containing 2% BSA. After washing, 50 µL of HRP-labelled cholera toxin B subunit (CTB-HRP; Fisher Scientific) at the indicated concentration was added and incubated for 2 h at room temperature (RT). After three final washes, 50 µL of 1-Step Ultra TMB-ELISA Substrate Solution (Fisher Scientific) was added. The reaction was stopped with 1 M H2SO4, and absorbance was measured at 450 nm.

### 3.3 Detection of loaded probiotics by ELISA

Bacteria at OD600 = 2, loaded with GM1 or GM1-cholesteryl-peptides in V-bottom plates, were washed three times with PBS containing 0.05% Tween 20 by centrifugation at 1000 × g for 10 min. The bacterial suspensions were then diluted in PBS containing 2% BSA and incubated for 1 h at RT to reduce nonspecific binding. Cells were washed once, resuspended in PBS containing 2% BSA, supplemented with 1 µg/mL HRP-labelled CTB (Fisher Scientific), and incubated for 2 h at 37°C with agitation. The cells were washed three times with PBS containing 0.05% Tween 20 and resuspended in PBS. Aliquots (50 µL) of the bacterial suspensions were transferred to an ELISA plate, TMB substrate was added, and the reaction was stopped with H2SO4 before absorbance was measured at 450 nm.

### 3.4 Stability assay

To assess retention of CTB-binding capacity after stress treatment, bacteria loaded with GM1 or GM1-cholesteryl-peptides were exposed to heating at 70°C for 15 min, freezing/thawing, lyophilization, or incubation at pH 2 for 1 h at 37°C before CTB-HRP detection as described above. The readout therefore reflects retention of CTB recognition of surface-accessible GM1 rather than direct quantification of membrane incorporation. No bacterial viability assay was performed after these treatments.

### 3.5 In vitro CTB-binding reduction assay using Vero cells (flow cytometry)

Vero cells (kidney epithelial cells derived from African green monkey; DSMZ) were cultured in McCoy’s medium (Gibco) supplemented with 10% FBS (Gibco) and 1% penicillin-streptomycin (Gibco), and seeded in 96-well cell-culture plates at 5 × 10^4 cells/well. Loaded bacteria were adjusted to OD600 = 2 in PBS containing 2% BSA and incubated for 1 h at RT. The bacterial cells were washed once by centrifugation at 1000 × g for 10 min, resuspended in PBS containing 2% BSA and 1 µg/mL Alexa Fluor 555-labelled CTB (CTB-555; Fisher Scientific), and incubated for 2 h at 37°C and 220 rpm. The suspensions were centrifuged at 1000 × g for 10 min, and the CTB-555-containing supernatants were transferred to Vero-cell monolayers after removal of the culture medium. Cells were incubated for 1 h at 37°C, after which 50 µL McCoy’s medium was added to each well and the cells were incubated overnight at 37°C before flow-cytometric acquisition on a MACSQuant Analyzer 16. Archived FCS files and FlowJo v10.8.1 workspaces were used to reconstruct the original analysis workflow. Events were first gated on FSC-A versus SSC-A using the population annotated as “live” in the original workspace. Because this gate was defined solely by scatter parameters, the term “live” is used here only as the historical gate label and does not indicate viability-dye-based exclusion of dead cells. Single cells were subsequently selected using FSC-A versus FSC-H, and mean B2-A (PE-A) fluorescence was quantified within the resulting population. The same polygon gate boundaries were applied across the two acquisition plates. Across the archived files, 3,073–17,774 total events per sample were recorded for Plate 1 and 3,973–13,122 for Plate 2. Figure 2 combines two separately recorded experimental datasets. Within each dataset, MFI values were normalized to the corresponding 0 µg/mL bacterial-loading condition (100% CTB-555 binding) before the datasets were combined. The available archival records did not permit retrospective confirmation that the two datasets originated from biologically independent bacterial preparations; accordingly, they are described as separately recorded experimental datasets rather than independent biological replicates. A representative gating strategy is shown in Appendix II, Supplementary Figure 5.

### 3.6 Statistical analysis

Replicate terminology. Technical replicates refer to repeated measurements of the same experimental sample within an assay, whereas experimental repeats or datasets refer to separately recorded assay runs. Where archival records did not establish biological independence, datasets are not described as biological replicates. Figure 1. Data are summarized as mean ± SD of the experimental repeats (n = 3 for GM1gs and GM1GL; n = 2 for GM1LL); the available archival documentation does not establish that these repeats represent biologically independent bacterial preparations. Stability assay (Figure 3). Technical duplicate measurements were averaged within each experimental repeat before summary. Data are presented as mean ± SD of the experimental repeats (n = 3 for GM1gs and GM1GL; n = 2 for GM1LL). Because of the limited number of experimental repeats, no inferential statistical testing was performed. Figure 2. Here, n = 2 denotes two separately recorded experimental datasets and should not be interpreted as confirmation of two biologically independent bacterial preparations. MFI values from the two datasets were normalized within each dataset to the corresponding 0 µg/mL loading condition (100% CTB-555 binding) and are presented as mean ± SD. Apparent IC50 values and 95% confidence intervals were estimated from both Y-value series by nonlinear least-squares fitting of a four-parameter logistic model, y = bottom + (top - bottom)/(1 + (x/IC50)^HillSlope), using Python 3.13 and SciPy 1.17.0. CTB-HRP sequestration assay (Appendix II, Supplementary Figure 4). In vitro CTB depletion by the loaded bacteria was quantified in Excel. A standard curve was generated by plotting absorbance at 450 nm against CTB-HRP concentration (µg/mL), allowing the concentration of unbound CTB-HRP in unknown samples to be estimated.

### 3.7 Use of generative AI-assisted tools

During revision of the manuscript, OpenAI ChatGPT (GPT-5.6 Sol; OpenAI; accessed October 2026) was used to assist with language editing, restructuring text in response to reviewer comments, preparation and harmonization of figure layouts and data visualizations, and development of analysis scripts applied to existing datasets, including calculation and presentation of apparent IC50 values and confidence intervals. No experimental data were generated by the AI tool. All scientific content, numerical results, analyses, interpretations, figures, and final wording were reviewed and verified by the authors.

## Supplementary Materials

The following supporting information is provided with this article: Appendix I, chemical synthesis and characterization of biotin-propargyl- and GM1-functionalized cholesteryl-peptides; Appendix II, optimization of loading conditions, ELISA-based detection and sequestration assays, and representative flow-cytometry gating for the Vero-cell assay.

## Author Contributions

Conceptualization, I.M.R. and S.G.; methodology, I.M.R., N.S.-H., O.B. and S.G.; investigation, I.M.R. and N.S.-H.; validation, I.M.R.; formal analysis, I.M.R.; data curation, I.M.R.; visualization, I.M.R.; writing - original draft preparation, I.M.R.; writing - review and editing, I.M.R., N.S.-H., O.B. and S.G.; supervision, N.S.-H., O.B. and S.G.; project administration, I.M.R. and S.G.; resources, O.B. and S.G.; funding acquisition, O.B. and S.G. All authors have read and agreed to the final version of the manuscript.

## Data Availability Statement

The data supporting the findings of this study, including source data for the revised figures and archived flow-cytometry files used for re-analysis, are included in the article/supplementary material where applicable and are available from the corresponding author upon reasonable request.

### Acknowledgments

The authors thank Dr. Lise Hafkenscheid for technical assistance with the Vero-cell flow-cytometry experiments. During the preparation and revision of this manuscript, the authors used OpenAI ChatGPT (GPT-5.6 Sol; OpenAI; accessed October 2026) to assist with language editing, manuscript restructuring, figure and data-visualization preparation, and analysis scripts applied to existing datasets. The authors reviewed and edited all AI-assisted output and take full responsibility for the content of this publication.

## References

1. Church, D. L. Major factors affecting the emergence and re-emergence of infectious diseases. Clin. Lab. Med. 24, 559–586 (2004).

2. O’Brien, S. J. & Halder, S. L. S. GI Epidemiology: Infection epidemiology and acute gastrointestinal infections. Aliment. Pharmacol. Ther. 25, 669–674 (2007).

3. Das, B., Verma, J., Kumar, P., Ghosh, A. & Ramamurthy, T. Antibiotic resistance in Vibrio cholerae: Understanding the ecology of resistance genes and mechanisms. Vaccine 38, A83–A92 (2020).

4. Tanwar, J., Das, S., Fatima, Z. & Hameed, S. Multidrug resistance: An emerging crisis. Interdiscip. Perspect. Infect. Dis. 2014, (2014).

5. Baptista, P. V. et al. Nano-strategies to fight multidrug resistant bacteria-”A Battle of the Titans”. Front. Microbiol. 9, 1–26 (2018).

6. Cummings RD, Schnaar RL, Esko JD, et al. Principles of Glycan Recognition. In: Varki A, Cummings RD, Esko JD, et al., editors. Essentials of Glycobiology [Internet]. 4th edition. Cold Spring Harbor (NY): Cold Spring Harbor Laboratory Press; 2022. Chapter 29.

7. Gallegos, K. M. et al. Shiga toxin binding to glycolipids and glycans. PLoS One 7, (2012).

8. Nishikawa, K. Recent progress of Shiga toxin neutralizer for treatment of infections by Shiga toxin-producing Escherichia coli. Arch. Immunol. Ther. Exp. (Warsz). 59, 239–247 (2011).

9. Takeda, T., Yoshino, K. I., Adachi, E., Sato, Y. & Yamagata, K. In vitro assessment of a chemically synthesized Shiga toxin receptor analog attached to Chromosorb P (Synsorb Pk) as a specific absorbing agent of Shiga toxin 1 and 2. Microbiol. Immunol. 43, 331–337 (1999).

10. Trachtman, H. et al. Effect of an Oral Shiga Toxin-Binding Agent on Diarrhea-Associated Hemolytic Uremic Syndrome in Children: A Randomized Controlled Trial. J. Am. Med. Assoc. 290, 1337–1344 (2003).

11. Das, S. et al. Neutralization of cholera toxin with nanoparticle decoys for treatment of cholera. PLoS Negl. Trop. Dis. 12, 1–17 (2018).

12. Lee, N. Y., Ko, W. C. & Hsueh, P. R. Nanoparticles in the treatment of infections caused by multidrug-resistant organisms. Front. Pharmacol. 10, 1–10 (2019).

13. Hsueh, B. Y. & Waters, C. M. Combating Cholera. F1000Research 8, F1000 Faculty Rev-589 (2019). doi:10.12688/f1000research.18093.1.

14. Duan, F. & March, J. C. Engineered bacterial communication prevents Vibrio cholerae virulence in an infant mouse model. Proc. Natl. Acad. Sci. U. S. A. 107, 11260–11264 (2010).

15. Mao, N., Cubillos-Ruiz, A., Cameron, D. E. & Collins, J. J. Probiotic strains detect and suppress cholera in mice. Sci. Transl. Med. 10, 1–9 (2018).

16. Zhou, Z. et al. Engineering probiotics as living diagnostics and therapeutics for improving human health. Microb. Cell Fact. 19, 1–12 (2020).

17. Paton, A. W., Morona, R. & Paton, J. C. Receptor-mimic probiotics: Potential therapeutics for bacterial toxin-mediated enteric diseases. Expert Rev. Gastroenterol. Hepatol. 4, 253–255 (2010).

18. Focareta, A., Paton, J. C., Morona, R., Cook, J. & Paton, A. W. A Recombinant Probiotic for Treatment and Prevention of Cholera. Gastroenterology 130, 1688–1695 (2006).

19. Allen, S. J., Martinez, E. G., Gregorio, G. V. & Dans, L. F. Probiotics for treating acute infectious diarrhoea. Cochrane Database Syst. Rev. 2010, (2010).

20. Patman, G. Diarrhoea: Lactobacillus reuteri reduces episodes of diarrhoea in healthy children. Nat. Rev. Gastroenterol. Hepatol. 11, 269 (2014).

21. Ahmadi, E., Alizadeh-Navaei, R. & Rezai, M. S. Efficacy of probiotic use in acute rotavirus diarrhea in children: A systematic review and meta-analysis. Casp. J. Intern. Med. 6, 187–195 (2015).

22. Guandalini, S. et al. Lactobacillus GG administered in oral rehydration solution to children with acute diarrhea: a multicenter European trial. J. Pediatr. Gastroenterol. Nutr. 30, 54–60 (2000). doi:10.1097/00005176-200001000-00018.

23. Aggarwal, S. et al. Lactobacillus GG for treatment of acute childhood diarrhoea: An open labelled, randomized controlled trial. Indian J. Med. Res. 139, 379–385 (2014).

24. Pant, A. R. et al. Lactobacillus GG and acute diarrhoea in young children in the tropics. J. Trop. Pediatr. 42, 162–165 (1996).

25. Sonnenborn, U. Escherichia coli strain Nissle 1917—from bench to bedside and back: history of a special Escherichia coli strain with probiotic properties. FEMS Microbiol. Lett. 363, fnw212 (2016). doi:10.1093/femsle/fnw212.

26. Mathipa, M. G. & Thantsha, M. S. Probiotic engineering: Towards development of robust probiotic strains with enhanced functional properties and for targeted control of enteric pathogens. Gut Pathog. 9, 1–17 (2017).

27. Bawa, A. S. & Anilakumar, K. R. Genetically modified foods: Safety, risks and public concerns -A review. J. Food Sci. Technol. 50, 1035–1046 (2013).

28. Heikkilä, J. E., Nybom, S. M. K., Salminen, S. J. & Meriluoto, J. A. O. Removal of cholera toxin from aqueous solution by probiotic bacteria. Pharmaceuticals 5, 665–673 (2012).

29. Tajima, G. et al. Impact of genetically modified organism requirements on gene therapy development in the EU, Japan, and the US. Mol. Ther. -Methods Clin. Dev. 26, 74–83 (2022).

30. Toutounian, K., Heinig, M. L., Götz, P. & Ulsemer, P. Exploring the pathogen binding potential within the human gut microbiome. Hum. Microbiome J. 18, 0–3 (2020).

31. Stuhr-Hansen, N. et al. Synthesis of Cholesterol-Substituted Glycopeptides for Tailor-Made Glycocalyxification of Artificial Membrane Systems. ChemBioChem 1403–1406 (2016) doi:10.1002/cbic.201600258.

32. Blixt, O. et al. Chemoenzymatic synthesis of 2-azidoethyl-ganglio-oligosaccharides GD3, GT3, GM2, GD2, GT2, GM1, and GD1a. Carbohydr. Res. 340, 1963–1972 (2005).

33. Vagianou, C. D. et al. ABO Blood Group Antigen Decorated Giant Unilamellar Vesicles Exhibit Distinct Interactions with Plasmodium falciparum Infected Red Blood Cells. ACS Chem. Biol. 13, 2421–2426 (2018).

34. Stuhr-Hansen, N., Vagianou, C. D. & Blixt, O. Synthesis of BODIPY-Labeled Cholesterylated Glycopeptides by Tandem Click Chemistry for Glycocalyxification of Giant Unilamellar Vesicles (GUVs). Chem. -A Eur. J. 23, 9472–9476 (2017).

35. Aureli, M. et al. GM1 Ganglioside: Past Studies and Future Potential. Mol. Neurobiol. 53, 1824–1842 (2016).

36. Zhang, H. et al. A pH-Responsive Gel Macrosphere Based on Sodium Alginate and Cellulose Nanofiber for Potential Intestinal Delivery of Probiotics. ACS Sustain. Chem. Eng. 6, 13924–13931 (2018).

