## Supplementary material for "In Vitro Assessment of Chemically Modified Probiotic Bacteria as Receptor-Mimic Decoys for Cholera Toxin B Subunit": IJMS_SUPPLEMENTARY_APPENDIX_I_ROUND1_FINAL.docx

**Supplementary Information-Appendix I**

**S1. Chemistry**

**S2. Synthesis of biotin-propargyl-functionalized cholesteryl-peptides**

*Biotin-OEG2-Pra-Lys-Lys-Ala(Cholesteryl)-Pro-OH*

*Biotin-OEG2-Pra-Glu-Lys-Ala(Cholesteryl)-Pro-OH*

**S3. Synthesis of biotin-GM1-functionalized cholesteryl-peptides by click chemistry**

*Biotin-OEG2-Ala(Triazyl-GM1)-Lys-Lys-Ala(Cholesteryl)-Pro-OH*

*Biotin-OEG2-Ala(Triazyl-GM1)-Glu-Lys-Ala(Cholesteryl)-Pro-OH*

**S1. Chemistry**

Unless otherwise stated, reagents and solvents were obtained from commercial suppliers and used without further purification. The GM1 azide used for click conjugation was obtained through the Consortium for Functional Glycomics (CFG) and used as received. HPLC was performed on a Dionex UltiMate 3000 system with UV detection using a Phenyl-Hexyl column (100 Å; Phenomenex). High-resolution mass spectrometry (HRMS) was performed on a Bruker Solarix XR ESI/MALDI-FT-ICR-MS instrument equipped with a 7 T magnet, and spectra were processed using Bruker DataAnalysis version 4.1.

**S2. Synthesis of biotin-propargyl-functionalized cholesteryl-peptides**

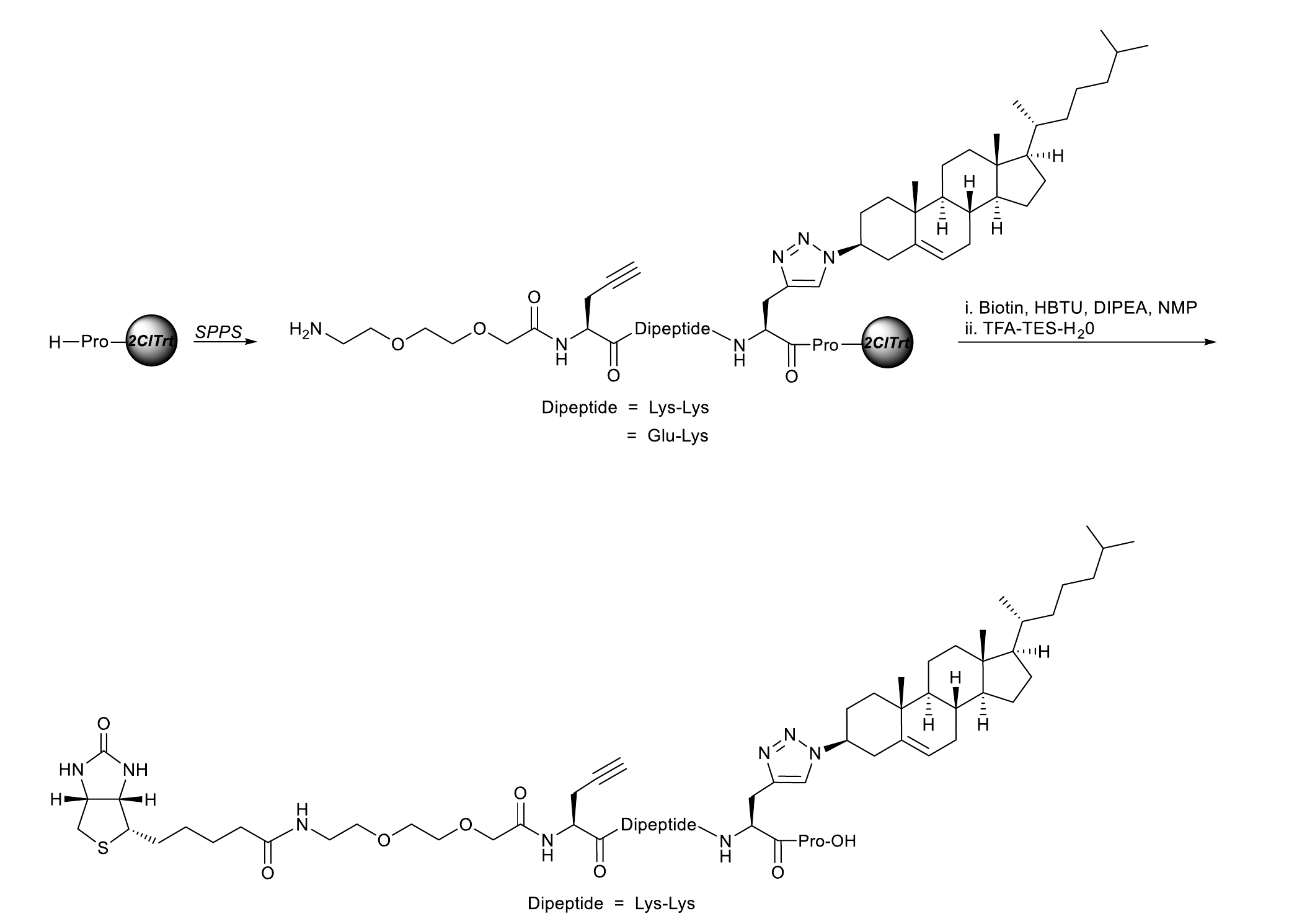

Lys-Lys- and Glu-Lys-spaced peptide precursors were synthesized on solid support by manual Fmoc-based solid-phase peptide synthesis (SPPS) using 2-chlorotrityl polystyrene resin (Sigma) preloaded with proline (loading, 1 mmol/g; 0.1 g, 0.1 mmol). Oligoethylene glycol, propargyl, and cholesteryl functionalities were introduced using Fmoc-OEG2-OH (Fmoc-O2Oc-OH; Iris), Fmoc-Pra-OH (Fmoc-propargyllysine; Iris), and Fmoc-Ala(Cholesteryl)-OH, respectively, as protected Fmoc amino-acid building blocks in the SPPS sequence. Each amino-acid coupling was performed with HBTU and DIPEA (resin/amino acid/HBTU/DIPEA, 1:4:4:8) in NMP (0.3 mL) for 30 min. After terminal Fmoc deprotection with 30% piperidine in DMF (1 × 5 min and 1 × 15 min; 1 mL for each treatment, with a DMF wash between treatments), the resin was washed with dichloromethane (5 × 4 mL) to afford the two corresponding resin-bound propargylated cholesteryl-peptides.

*Biotin-OEG2-Pra-Lys-Lys-Ala(Cholesteryl)-Pro-OH*

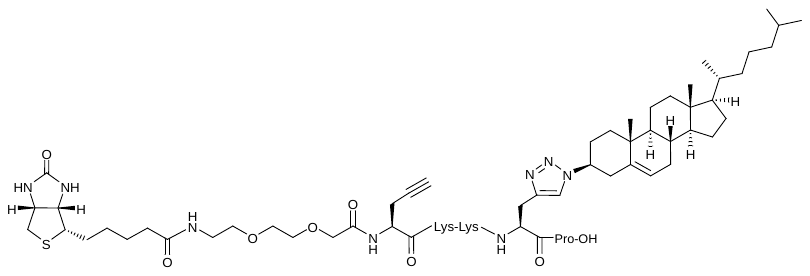

Biotinylation was performed by treating one tenth of the Lys-Lys-dipeptide-spaced resin with a preactivated solution of biotin (12 mg, 0.05 mmol) and HBTU (19 mg, 0.05 mmol) in DIPEA/NMP (1:6, 0.5 mL) for 30 min. After draining, the resin was washed with DMF (4×) followed by dichloromethane (5×). A TFA/triethylsilane (TES)/H2O cleavage cocktail (18:1:1, 1 mL) was then added and the resin was shaken for 1 h. The cleavage solution was evaporated, acetonitrile/water (1:1, 1 mL) was added, and the mixture was stirred vigorously for 2 min and filtered to obtain a clear solution. HPLC purification (Phenyl-Hexyl column, 100 Å; Phenomenex) using a water/acetonitrile gradient containing 0.1% formic acid, followed by lyophilization, yielded the biotin-terminated Lys-Lys-spaced propargylated cholesteryl-peptide (3.7 mg, 28%) as a white, fluffy powder. Purity was >98% by HPLC. HRMS (ES+) calcd for C78H115N13O11S [M + 2H+]2+, 672.9280; found, m/z 672.9265.

**HPLC purification (254 nm):**

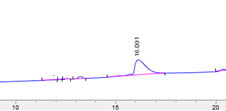

**MALDI:**

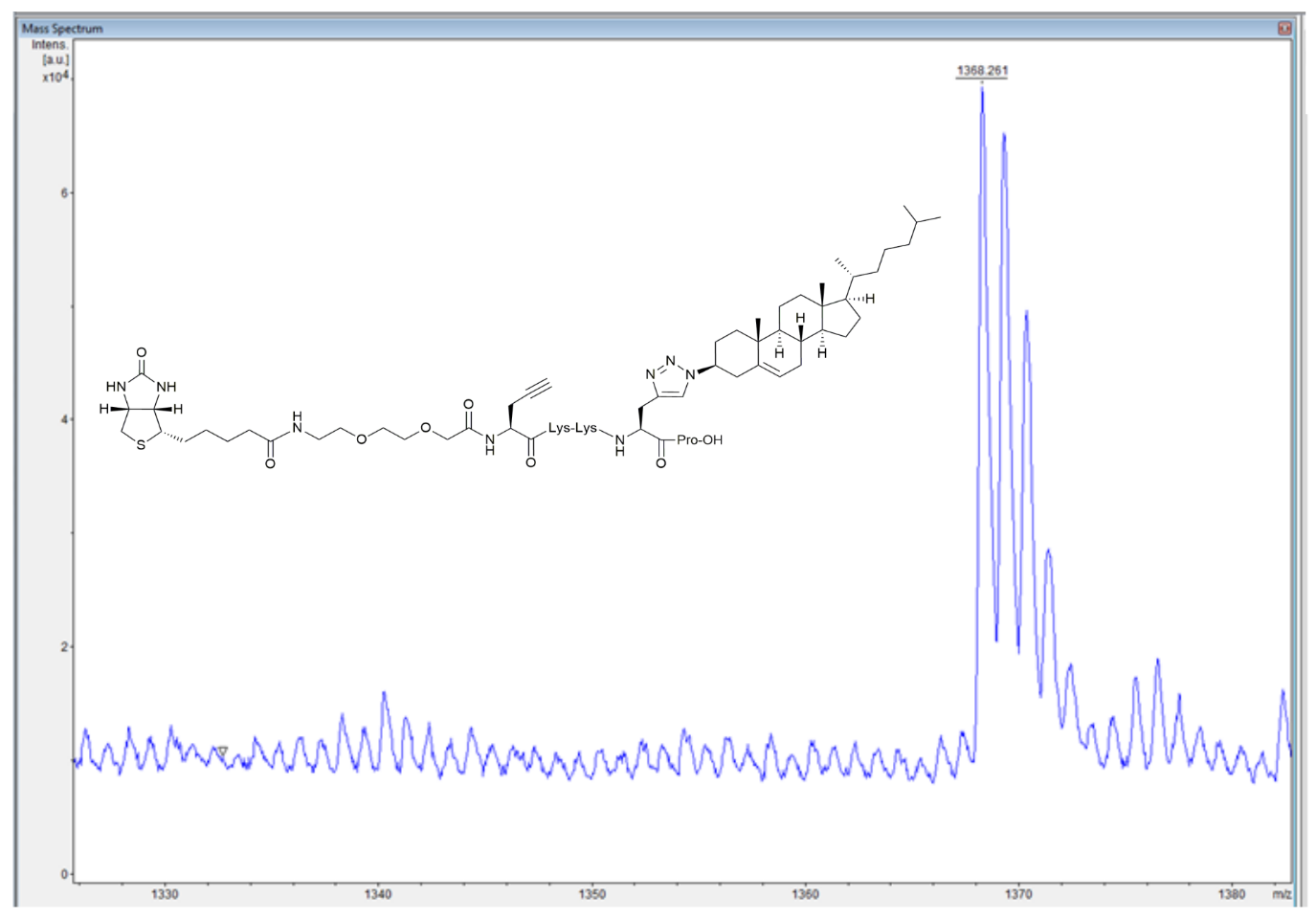

*Biotin-OEG2-Pra-Glu-Lys-Ala(Cholesteryl)-Pro-OH*

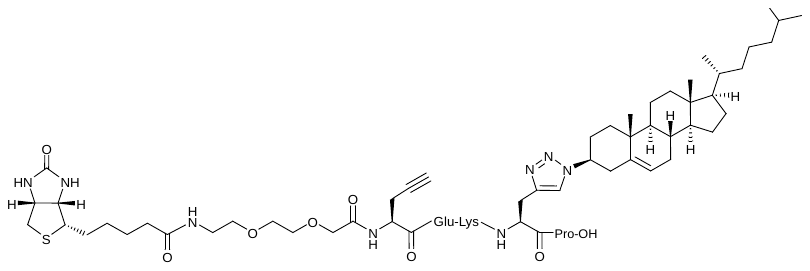

Biotinylation was performed by treating one tenth of the Glu-Lys-dipeptide-spaced resin with a preactivated solution of biotin (12 mg, 0.05 mmol; Iris) and HBTU (19 mg, 0.05 mmol) in DIPEA/NMP (1:6, 0.5 mL) for 30 min. After draining, the resin was washed with DMF (4×) followed by dichloromethane (5×). A TFA/triethylsilane (TES)/H2O cleavage cocktail (18:1:1, 1 mL) was then added and the resin was shaken for 1 h. The cleavage solution was evaporated, acetonitrile/water (1:1, 1 mL) was added, and the mixture was stirred vigorously for 2 min and filtered to obtain a clear solution. HPLC purification (Phenyl-Hexyl column, 100 Å; Phenomenex) using a water/acetonitrile gradient containing 0.1% formic acid, followed by lyophilization, yielded the biotin-terminated Glu-Lys-spaced propargylated cholesteryl-peptide (4.1 mg, 31%) as a white, fluffy powder. Purity was >98% by HPLC. HRMS (ES+) calcd for C69H110N12O13S [M + 2H+]2+, 673.4018; found, m/z 673.4006.

**HPLC purification (254 nm):**

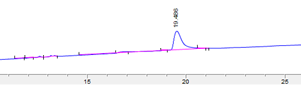

**MALDI:**

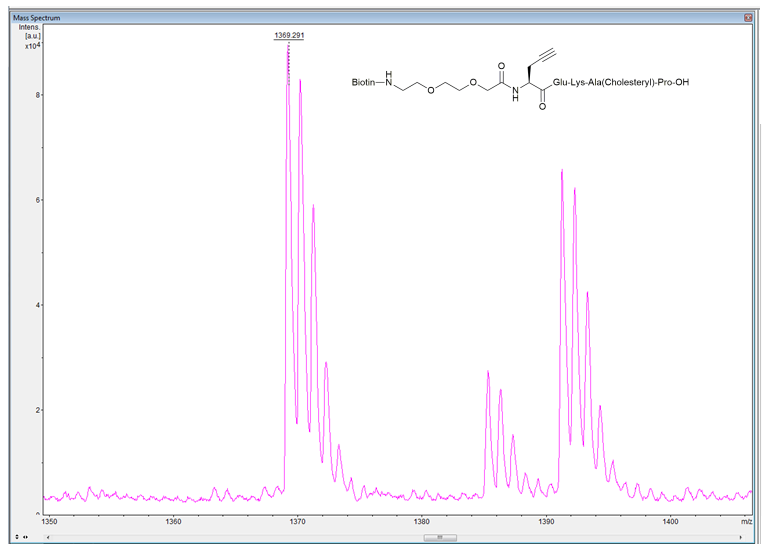

**ESI:**

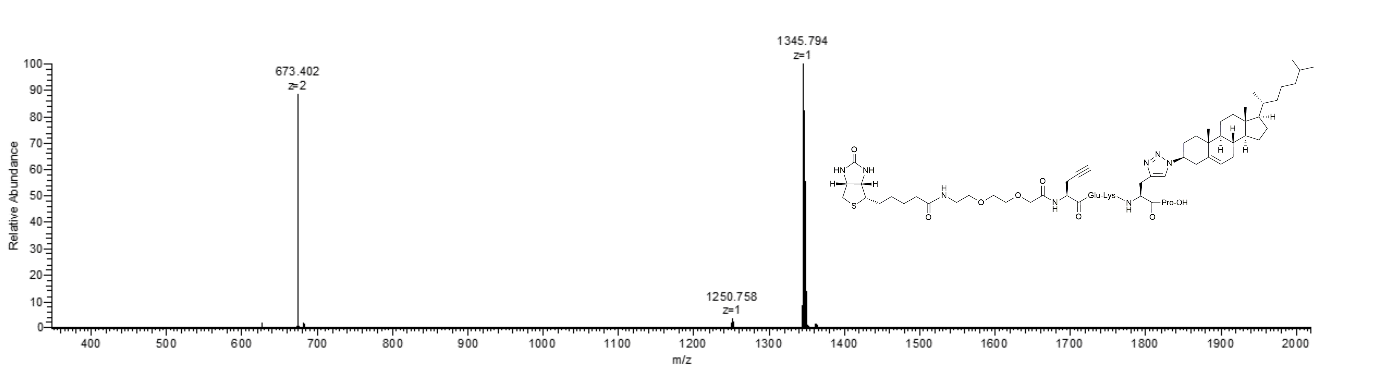

**S3. Synthesis of biotin-GM1-functionalized cholesteryl-peptides by click chemistry**

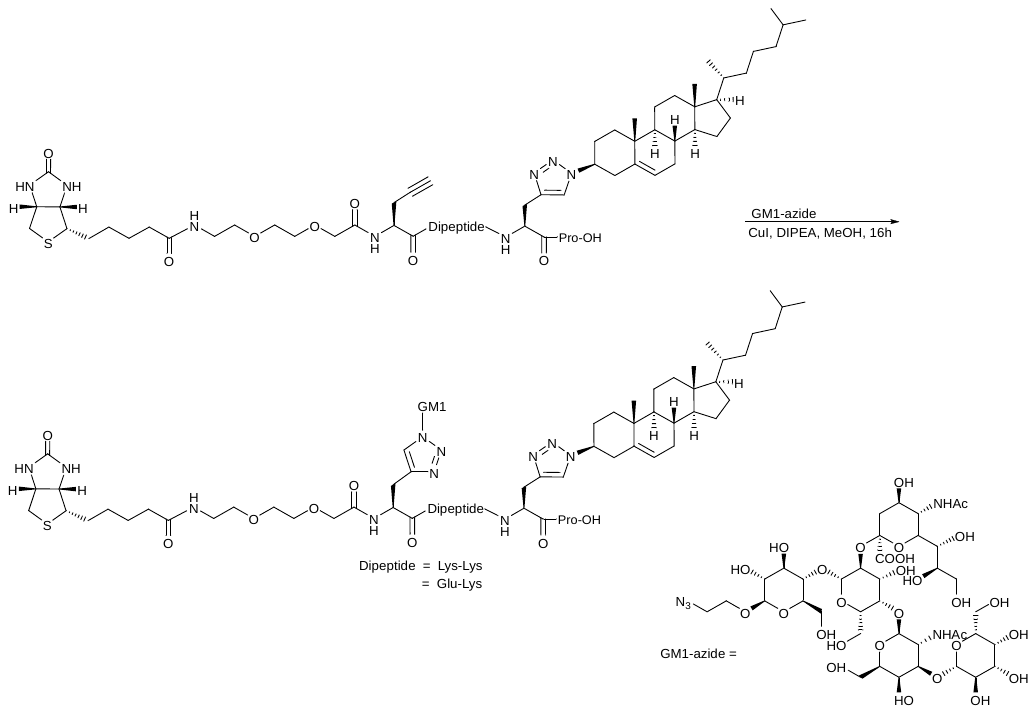

Biotin-terminated Lys-Lys- or Glu-Lys-spaced propargylated cholesteryl-peptide (1.1 mg, 0.3 μmol) and GM1 azide (0.33 mg, 0.31 μmol; obtained through the Consortium for Functional Glycomics, CFG) were dissolved in thoroughly argon-degassed DIPEA/methanol (1:9, 0.1 mL) and added to CuI (0.0019 mg, 0.01 μmol), prepared from a nitrogen-evaporated aliquot of a vigorously stirred copper-salt solution in methanol. The reaction mixture was shaken at 500 rpm for 16 h and then evaporated to dryness. The resulting pale azure-blue oily residue was redissolved in 1% formic acid in H2O/MeCN (1:1, 1 mL), filtered, and lyophilized.

*Biotin-OEG2-Ala(Triazyl-GM1)-Lys-Lys-Ala(Cholesteryl)-Pro-OH*

The product was obtained in quantitative yield as a colorless oil. Analytical purification, which substantially reduced the isolated yield, was performed by Sep-Pak C18 column filtration using an H2O/MeCN (3:2) gradient. HRMS (ES+) calcd for C109H180N18O40S [M + 2H+]2+, 1206.6162; found, m/z 1206.6124.

**MALDI:**

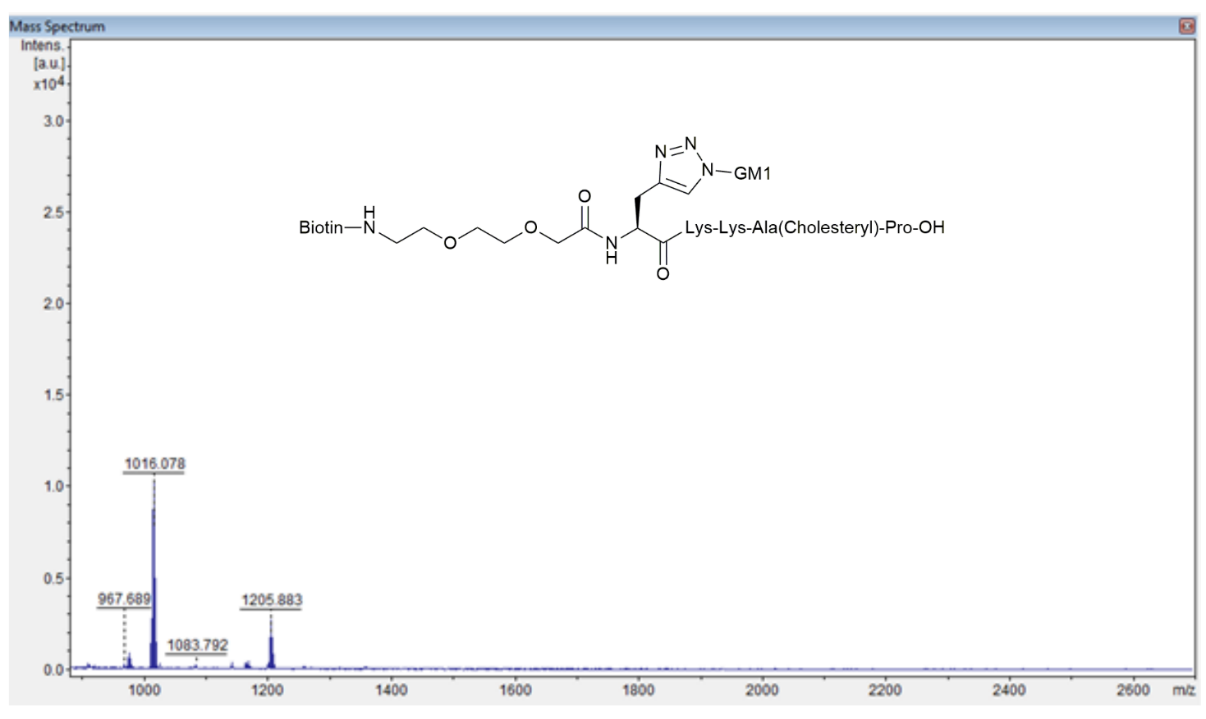

*Biotin-OEG2-Ala(Triazyl-GM1)-Glu-Lys-Ala(Cholesteryl)-Pro-OH*

The product was obtained in quantitative yield as a colorless oil. Analytical purification, which substantially reduced the isolated yield, was performed by Sep-Pak C18 column filtration using an H2O/MeCN (3:2) gradient. HRMS (ES+) calcd for C108H175N17O42S [M + 2H+]2+, 1207.0901; found, m/z 1207.0860.

**MALDI:**

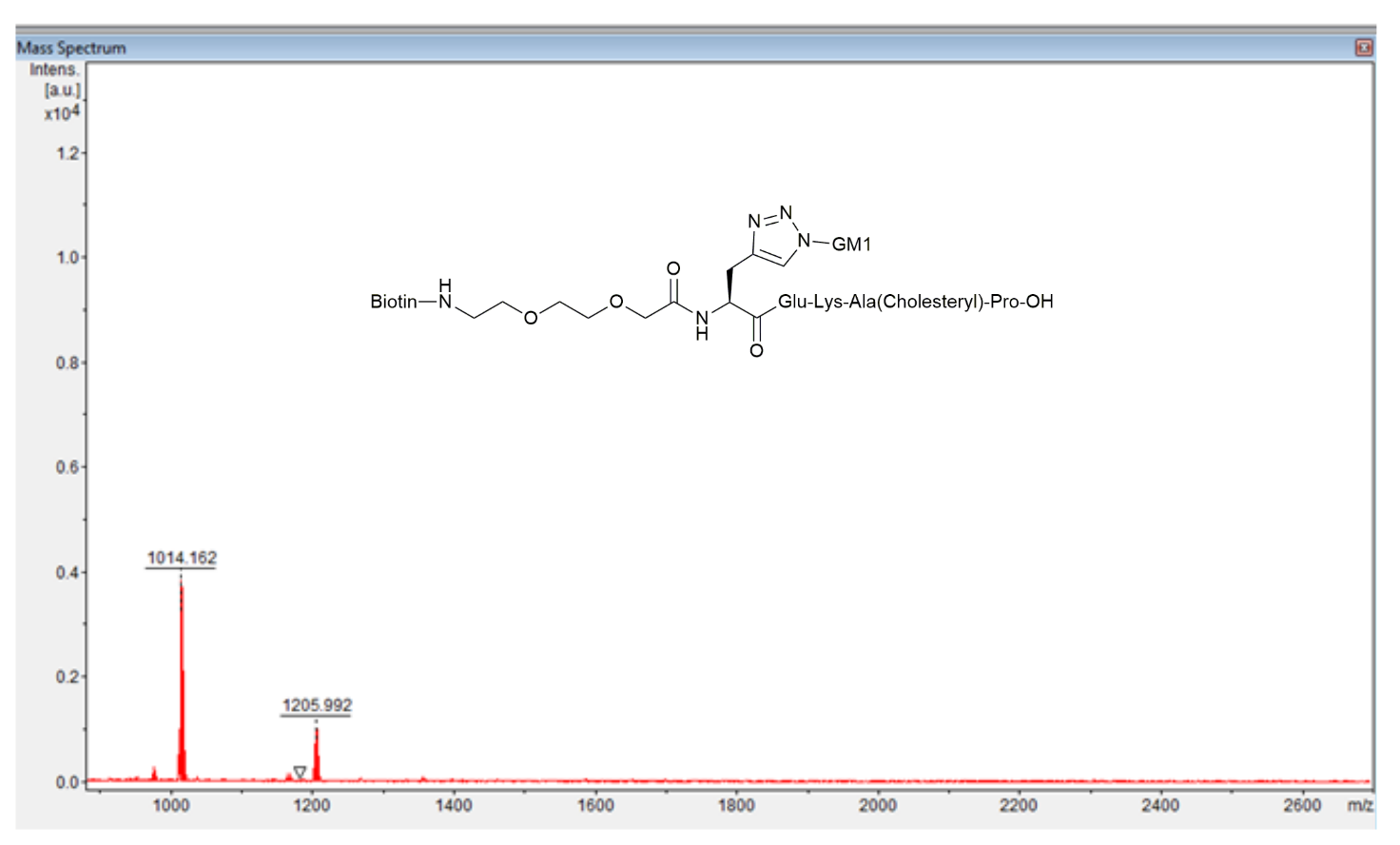
