## Supplementary material for "In Vitro Assessment of Chemically Modified Probiotic Bacteria as Receptor-Mimic Decoys for Cholera Toxin B Subunit": IJMS_SUPPLEMENTARY_APPENDIX_II_ROUND1_FINAL.docx

**Supplementary Information-Appendix II**

**S1. Optimization of loading conditions**

**S2. Detection of loaded probiotics at different OD600 values (ELISA)**

**S3. In vitro CTB-HRP sequestration assay for loaded probiotics (ELISA)**

**S4. Representative flow-cytometry gating strategy for the Vero-cell assay**

S1. Optimization of loading conditions

To investigate the loading conditions associated with maximal CTB-HRP binding/sequestration, we used natural GM1 ganglioside as an initial reference and assessed a range of bacterial cell densities (OD600 0.5-7) and GM1 loading concentrations (0-10 µg/mL), as described in the Methods section (Preparation of loaded probiotics). After removal of unbound GM1 by three washes in PBS + 0.05% Tween 20, bacteria were incubated with 1 µg/mL CTB-HRP for 2 h at 37°C, 220 rpm. Residual unbound CTB-HRP in the supernatant was transferred to GM1-coated ELISA plates and detected by TMB/H2SO4 development (Sup. Figure 1A). For both LR and EcN, GM1 loading concentrations of approximately 1-2.5 µg/mL corresponded to the plateau region associated with maximal depletion of 1 µg/mL CTB-HRP from the supernatant in this assay (Sup. Figure 1B,C). These single-experiment data identify an assay-specific concentration range associated with maximal CTB-HRP depletion but do not establish stoichiometric binding or neutralization of intact cholera toxin. Lower bacterial densities showed a modestly greater reduction in free CTB-HRP signal (Sup. Figure 1B,C).

Based on Sup. Figure 1, we next evaluated the highest CTB-HRP concentration that could be depleted from the supernatant by GM1-loaded bacteria. Bacterial densities OD600 0.5-7 were loaded with a fixed 2.5 µg/mL concentration of GM1 and exposed to increasing concentrations of CTB-HRP (0-2.5 µg/mL) (Sup. Figure 2A). For LR, 2.5 µg/mL GM1 loading was associated with strong depletion of CTB-HRP across the tested concentration range, with no clear dependence on bacterial density (Sup. Figure 2B). No clear correlation between OD600 and the residual CTB-HRP signal was observed for EcN (Sup. Figure 2C). Because these measurements derive from a single experiment and include washing and transfer steps, they are interpreted as assay-specific CTB-HRP sequestration readouts rather than quantitative stoichiometry or evidence of intact holotoxin neutralization.

S2. Detection of loaded probiotics at different OD600 values (ELISA)

Bacteria at OD600 = 2, 4, or 6 were loaded with GM1, GM1-cholesteryl-peptides, or the corresponding controls in V-bottom plates and washed three times with PBS containing 0.05% Tween 20, as described in the Methods section. The cells were diluted in 200 μL/well PBS containing 2% BSA and incubated at room temperature (RT) for 1 h to reduce nonspecific binding. After one wash, the cells were resuspended in PBS containing 2% BSA and 1 μg/mL CTB-HRP and incubated for 2 h at 37°C with shaking at 220 rpm. Cells were washed three times with PBS containing 0.05% Tween 20 and resuspended in 50 μL/well PBS. The bacterial suspensions were transferred from the deep V-bottom polypropylene plate to an ELISA plate. TMB substrate solution (50 μL/well) was added, the reaction was stopped with 50 μL/well 1 M H2SO4, and absorbance was measured at 450 nm (Sup. Figure 3A).

S3. In vitro CTB-HRP sequestration assay for loaded probiotics (ELISA)

Microorganisms loaded with GM1 or GM1-cholesteryl-peptides as described above were adjusted to OD600 = 2 in PBS containing 2% BSA and incubated at RT for 1 h. The cells were washed once, resuspended in PBS containing 2% BSA and 1 μg/mL CTB-HRP, and incubated for 2 h at 37°C with shaking at 220 rpm. The bacterial suspensions were centrifuged, and supernatants containing residual unbound CTB-HRP were transferred to Maxisorp 96-well ELISA plates coated with 1 μg/mL GM1 and incubated for 2 h at RT. CTB-HRP that had not been preincubated with loaded bacteria was used to generate a standard curve relating absorbance at 450 nm to CTB-HRP concentration. Comparison of the supernatant signal with this standard curve allowed estimation of residual unbound CTB-HRP and, consequently, apparent in vitro CTB-HRP sequestration by the GM1-lipid-loaded microorganisms (Sup. Figure 4).

**S4. Representative flow-cytometry gating strategy for the Vero-cell assay**

For the archived Vero-cell acquisition used to reconstruct the Figure 2 analysis workflow, original FCS files and FlowJo v10.8.1 workspaces were examined. Events were first selected on FSC-A versus SSC-A using the polygon population annotated as “live” in the original workspace. Because this gate was defined only by scatter parameters, the historical label “live” is not interpreted as evidence of viability-dye-based dead-cell exclusion. Single-cell events were subsequently selected using FSC-A versus FSC-H, and mean B2-A (PE-A) fluorescence was quantified within the resulting “live/Single Cells” population. The same polygon gate boundaries were applied across both plates of this archived acquisition.

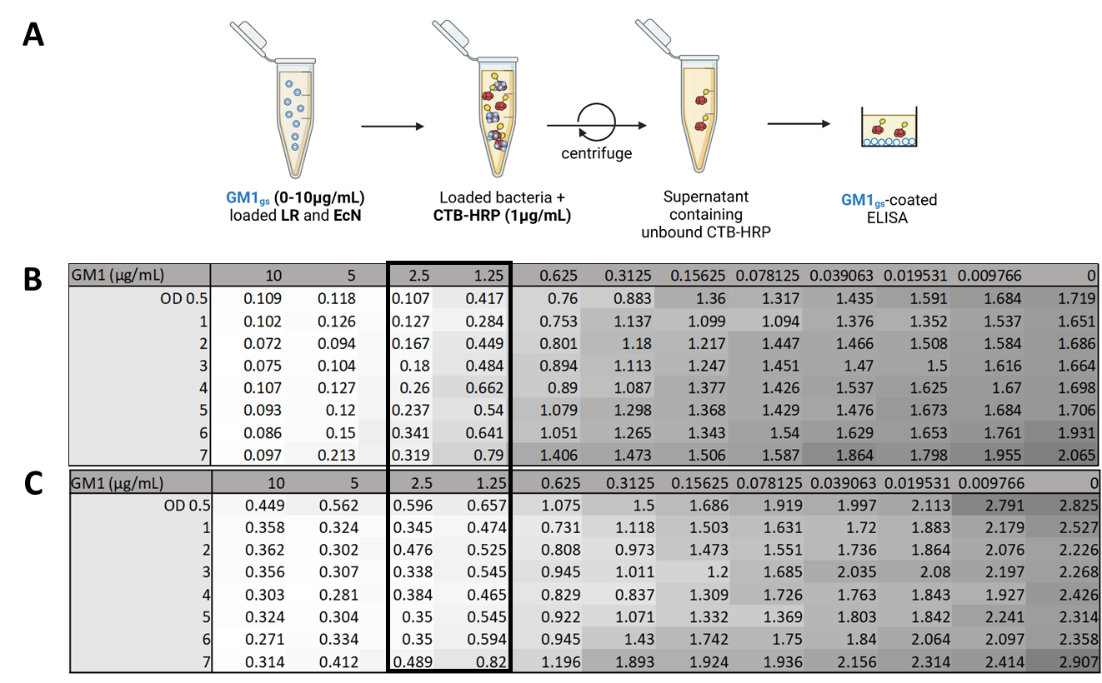

**Sup. Figure 1.** Experimental setup (A). Residual CTB-HRP signal after preincubation of 1 μg/mL CTB-HRP with LR and EcN loaded with 0–10 μg/mL GM1 (B and C, respectively). The black rectangle indicates the GM1 concentration range corresponding to the plateau of CTB-HRP depletion in this assay. Data are from a single experiment.

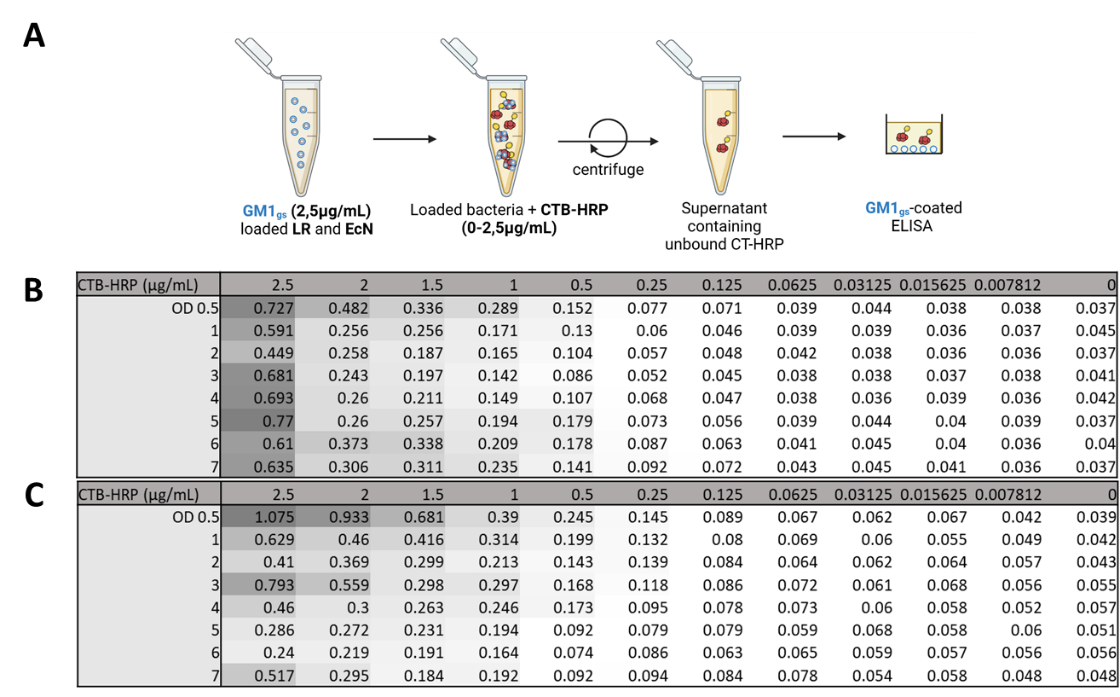

**Sup. Figure 2.** Experimental setup (A). Residual CTB-HRP signal after preincubation of 0–2.5 μg/mL CTB-HRP with LR and EcN loaded with 2.5 μg/mL GM1 (B and C, respectively). Data are from a single experiment.

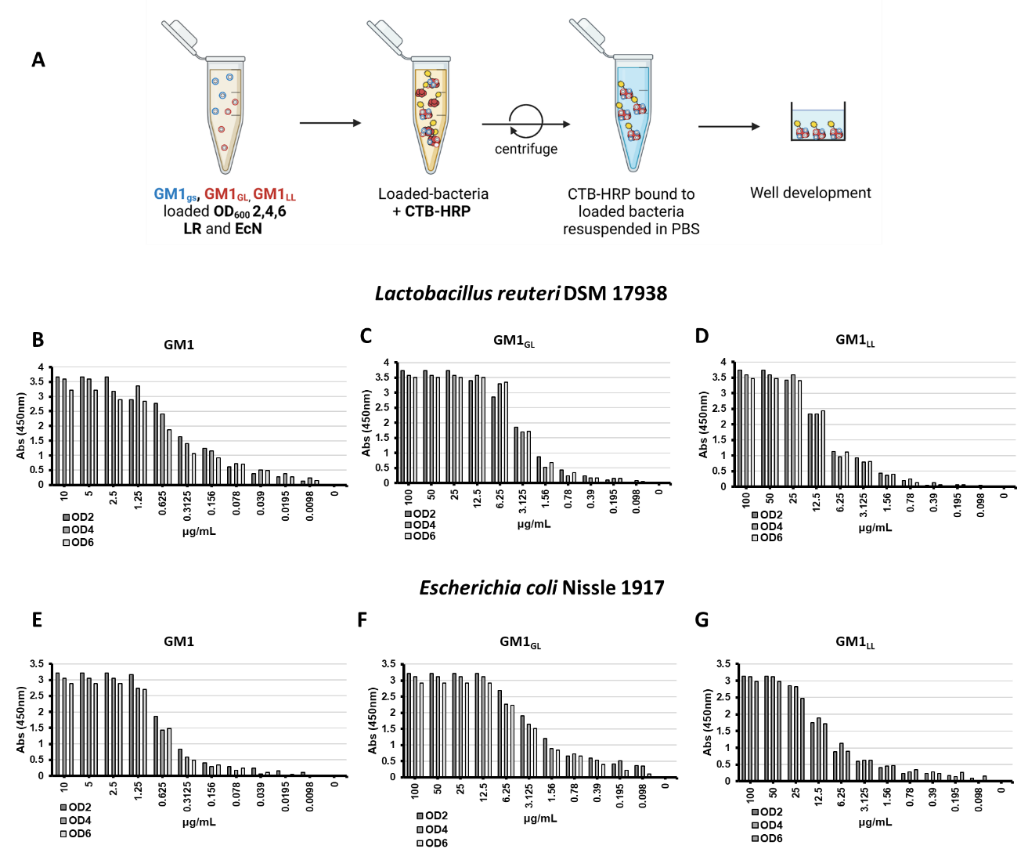

**Sup. Figure 3.** CTB-HRP binding to GM1-loaded LR and EcN at different bacterial densities (OD600 = 2, 4, and 6). A) Experimental setup. CTB-HRP binding to LR loaded with GM1gs (B), GM1GL (C), or GM1LL (D), and to EcN loaded with GM1gs (E), GM1GL (F), or GM1LL (G). CTB-HRP recognition is used here as an indirect readout of surface-accessible GM1 rather than as a direct measurement of membrane incorporation.

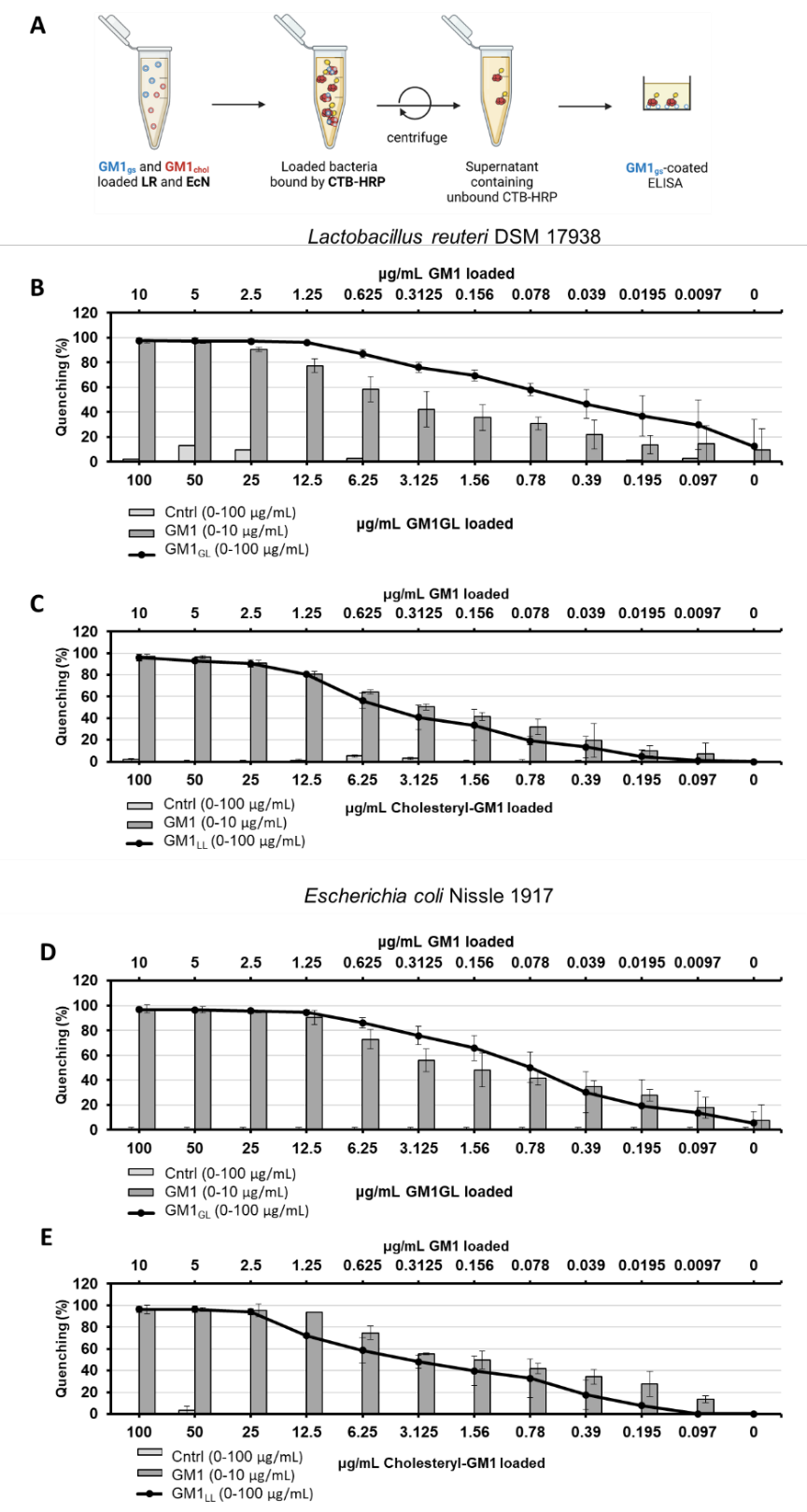

**Sup. Figure 4.** Concentration-dependent sequestration of CTB-HRP by GM1-lipid-loaded LR and EcN. A) Graphical description of the assay. OD600 = 2 LR and EcN loaded with increasing concentrations of GM1gs (0–10 μg/mL), GM1-cholesteryl-peptides, or the corresponding cholesteryl controls (0–100 μg/mL) were incubated with CTB-HRP. After centrifugation, residual unbound CTB-HRP in the supernatant was transferred to a GM1-coated ELISA plate (1 μg/mL). B) Apparent CTB-HRP sequestration by GM1GL-loaded LR. C) Apparent CTB-HRP sequestration by GM1LL-loaded LR. D) Apparent CTB-HRP sequestration by GM1GL-loaded EcN. E) Apparent CTB-HRP sequestration by GM1LL-loaded EcN. The GM1gs concentration-response curve (0–10 μg/mL) is shown in the corresponding panels for comparison.

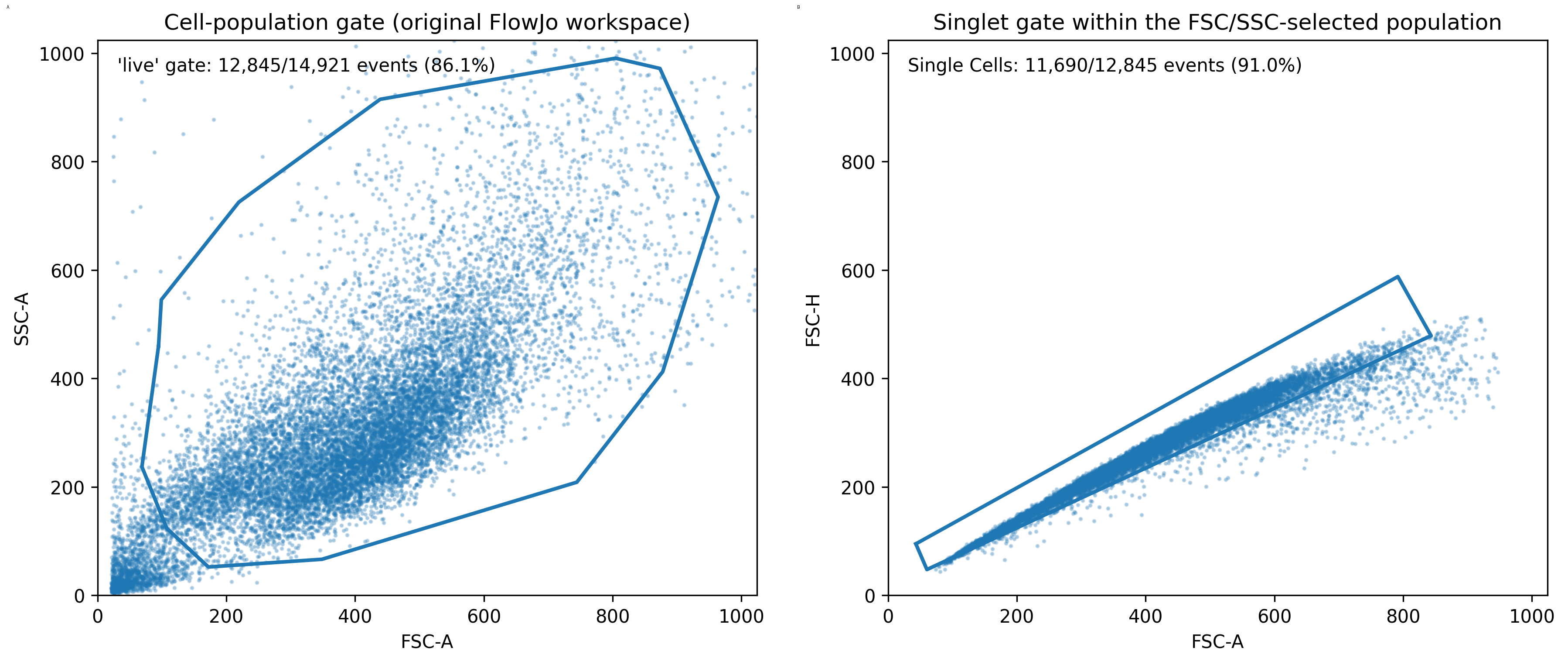

**Sup. Figure 5.** Representative flow-cytometry gating strategy for the archived Vero-cell acquisition. A) FSC-A versus SSC-A polygon gate corresponding to the population annotated as “live” in the original FlowJo workspace. The representative sample shown contains 14,921 total events, of which 12,845 fall within this scatter-defined gate. B) FSC-A versus FSC-H singlet gate applied within the FSC/SSC-selected population; 11,690 events fall within the Single Cells gate in the representative sample. The same polygon gate boundaries were applied across both plates of this archived acquisition. The historical “live” label reflects the original workspace nomenclature and does not indicate viability-dye-based discrimination.
